# Evolution and codon usage bias of mitochondrial and nuclear genomes in *Aspergillus* section *Flavi*

**DOI:** 10.1101/2022.08.23.505037

**Authors:** Miya Hugaboom, E. Anne Hatmaker, Abigail L. LaBella, Antonis Rokas

**Affiliations:** Department of Biological Sciences, Vanderbilt University, Nashville, TN, USA; Evolutionary Studies Initiative, Vanderbilt University, Nashville, TN, USA; Department of Bioinformatics and Genomics, University of North Carolina at Charlotte, Charlotte, NC USA

**Keywords:** mitogenome, phylogeny, codon usage bias, *Aspergillus*, section *Flavi*, fungi

## Abstract

The fungal genus *Aspergillus* contains a diversity of species divided into taxonomic sections of closely related species. Section *Flavi* contains 33 species, many of industrial, agricultural, or medical relevance. Here, we analyze the mitochondrial genomes (mitogenomes) of 20 *Flavi* species—including 18 newly assembled mitogenomes—and compare their evolutionary history and codon usage bias (CUB) patterns to their nuclear counterparts. CUB refers to variable frequencies of synonymous codons in coding DNA and is shaped by a balance of neutral processes and natural selection. All mitogenomes were circular DNA molecules with highly conserved gene content and order. As expected, genomic content, including GC content, and genome size differed greatly between mitochondrial and nuclear genomes. Phylogenetic analysis based on 14 concatenated mitochondrial genes predicted evolutionary relationships largely consistent with those predicted by a phylogeny constructed from 2,422 nuclear genes. Comparing similarities in interspecies patterns of CUB between mitochondrial and nuclear genomes showed that species grouped differently by patterns of CUB depending on whether analyses were performed using mitochondrial or nuclear relative synonymous usage values. We found that patterns of CUB at gene-level are more similar between mitogenomes of different species than the mitogenome and nuclear genome of the same species. Finally, we inferred that, although most genes—both nuclear and mitochondrial—deviated from the neutral expectation for codon usage, mitogenomes were not under translational selection while nuclear genomes were under moderate translational selection. These results contribute to the study of mitochondrial genome evolution in filamentous fungi.

## Introduction

The fungal genus *Aspergillus* is an important genus of filamentous fungi. The genus houses species with industrial applications, important pathogens of humans, animals and crops, producers of potent carcinogenic mycotoxins, and the genetic model organism *Aspergillus nidulans* (de Vries et al. 2017). *Aspergillus* is divided into taxonomic sections of closely related species. Section *Flavi* consists of 33 species, many of which have industrial, agricultural, or medical relevance (Gourama and Bullerman 1995; Hedayati et al. 2007; de Vries et al. 2017; Frisvad et al. 2018; Homa et al. 2019). For example*, A. oryzae* constitutes an important cell factory for enzyme production and, along with *A. sojae*, is vital to the production of a range of fermented foods (Machida et al. 2008; Sato et al. 2011). Conversely*, A. flavus* is an effective producer of aflatoxin B, a potent carcinogenic mycotoxin, and has been found to be both a plant contaminant and occasional pathogen, as well as anopportunistic human pathogen (Hedayati et al. 2007; Hoffmeister and Keller 2007, Dolezal et al. 2014). To better understand the diversity of these fungi, a recent study sequenced the genomes for 23 of the 33 known *Flavi* species to gain insights into their biology and evolution (Kjærbølling et al. 2020).

Previous genomic analyses of section *Flavi* focus almost exclusively on the nuclear genomes of the sequenced species (de Vries et al. 2017; Kjærbølling et al. 2020); the sole exception was a 2012 study that described the genomes of six diverse *Aspergillus* species, including two from section *Flavi* (Joardar et al. 2012). However, whole genome sequencing captures nucleotide sequences from both nuclear and organellar genomes. Fungal mitochondria have been linked to diverse processes including energy metabolism, cell differentiation, drug resistance, biofilm and hyphal growth regulation, and virulence, amongst others (Sanglard et al. 2001; Burger et al. 2003; Martins et al. 2011; Chatre and Ricchetti 2014; Calderone et al. 2015). Using appropriate software, mitochondrial reads can be effectively filtered and separated from nuclear reads within existing whole-genome sequencing datasets to be used for mitochondrial genome (mitogenome) assembly and annotation (Hugaboom et al. 2021). Fungal mitogenomes, including those of *Aspergillus* species, are typically circular and composed of a single chromosome (Joardar et al. 2012). Mitogenomes replicate independently from the nuclear genome and cell cycle and tend to have high copy number. Fourteen protein-coding genes involved in the electron transport chain are highly conserved within fungal mitogenomes (Gray et al. 1999; Lavín et al. 2008; Joardar et al. 2012). Genes for two ribosomal rRNAs subunits, one large and one small, and a variable number of tRNAs also tend to be housed in the mitogenome (Gray et al. 1999; Lavín et al. 2008; Joardar et al. 2012). Variation in fungal mitogenomes is largely due to differences in intron distribution and the variable presence of accessory mitochondrial genes, even between closely related species (Joardar et al. 2012; Li, Xiang, et al. 2019; Li, Wang, et al. 2019; Wang et al. 2020; Zhang et al. 2020; Chen et al. 2021). Importantly, mitogenomes also differ from nuclear genomes in their inheritance pattern. Whereas fungal nuclear genomes are inherited from both parents regardless of mating type and display genetic recombination, fungal mitogenomes are uniparentally inherited and rarely display recombination, offering a unique phylogenetic perspective (Santamaria et al. 2009; Kjærbølling et al. 2020). Mitochondrial genomes may therefore hold clues to both the biology and evolution of these fungal species.

Another key difference between mitogenomes and nuclear genomes is codon usage bias (CUB). CUB refers to the different frequency of synonymous codons—those that code for the same amino acid—in coding DNA. Changes in synonymous codons do not alter primary protein sequence and were thus once assumed to be selectively neutral (Jia and Higgs 2008; Wei et al. 2014; LaBella et al. 2019). However, CUB has been found to influence numerous cellular processes, particularly those associated with translation (Stoletzki and Eyre-Walker 2007; Zhou et al. 2009). This is hypothesized to be due to codon optimization: the tendency for codon usage to be correlated to the abundance of tRNA molecules in the genome (Post et al. 1979; Nakamura et al. 1980; Ikemura 1981; Gouy and Gautier 1982; P M Sharp and Li 1986; Thomas et al. 1988). During translation, mRNAs containing optimized codons—codons corresponding to the tRNA pool of the cell—are translated more efficiently than those with non-optimal codon usage (Bulmer 1991; Xia 1998; Chevance et al. 2014; Presnyak et al. 2015; Hanson and Coller 2018). In many organisms, this leads to a correlation between codon usage and protein production (Ikemura 1981; Bulmer 1991; Gustafsson et al. 2004; Hiraoka et al. 2009; Roymondal et al. 2009; Zhipeng et al. 2016; Payne and Alvarez-Ponce 2019; Sahoo et al. 2019). Importantly, mitogenomes house their own set of tRNAs that is distinct from that of the nuclear genome and thus may exhibit patterns of CUB shaped by optimization to a greater extent by the mitochondrial set of tRNAs (tRNAome) than the nuclear tRNAome. Variation in synonymous codon usage is a widespread phenomenon at codon, gene, and whole genome levels in nuclear and mitochondrial genomes (LaBella et al. 2019; LaBella et al. 2021; Wint et al. 2022). This variation in codon usage likely reflects a balance of mutational bias (e.g., GC content), natural selection (e.g., translational selection), and genetic drift (Ikemura 1985; Shields and Sharp 1987; Sharp et al. 1993; Wei et al. 2014). The balance of these forces varies between organisms. In many microbes, for example, translational selection plays a large role, whereas mutational bias plays the primary role in humans (Sharp et al. 1993). However, analysis of mitochondrial CUB in fungi is limited (Kamatani and Yamamoto 2007; Carullo and Xia 2008). Understanding patterns of CUB can provide insight into the evolutionary history of individual genes and entire genomes.

To gain insights into the evolution of mitogenomes from the section *Flavi,* we analyzed mitochondrial genomes of 20 section *Flavi* species—including 18 newly assembled ones—and compared their phylogeny and CUB to the nuclear genomes of the same species. All mitogenomes were confirmed to be circular DNA molecules of low GC content with highly conserved gene content and gene order. Genomic content and size differed greatly between mitochondrial and nuclear genomes. We then inferred and compared phylogenies constructed from mitochondrial versus nuclear genes. The widespread presence and high copy number of mitogenomes within the cell as well as the apparent lack of recombination make mitochondrial genes and genomes useful markers for phylogenetic analyses. Currently, phylogenies constructed for *Aspergillus* section *Flavi* are based solely on nuclear genome markers (Kjærbølling et al. 2020; Shen et al. 2020). Phylogenetic analysis based on 14 concatenated mitochondrial genes (mitogenes) predicted evolutionary relationships largely consistent with those inferred by a phylogeny based on nuclear data. We then investigated CUB in mitochondrial and nuclear genomes. At the gene-level, we found that patterns of CUB reflect whether the gene is mitochondrial or nuclear in origin as well as mitogene identity rather than species of origin; these patterns were influenced largely by GC content of the third codon position. Finally, we determined that although most genes—both nuclear and mitochondrial—deviated from the neutral expectation, mitogenomes were not under translational selection while nuclear genomes were under moderate translational selection. By providing mitogenome assemblies for 20 section *Flavi* species and comparing the evolution of mitochondrial and nuclear genes in section *Flavi*, our study advances our understanding of genome evolution in the genus *Aspergillus*.

## Methods

### Genomic Data

We used strains from 21 species within section *Flavi* and *Aspergillus niger* (section *Nigri*) as an outgroup for phylogenetic analyses. For the mitochondrial dataset, we used a combination of available mitochondrial reference genomes and newly assembled whole-genome sequencing reads. Three previously assembled mitochondrial reference genomes (*Aspergillus sojae*, *Aspergillus oryzae,* and *Aspergillus niger*) were downloaded from NCBI’s Nucleotide Database (Juhász et al. 2008; Machida et al. 2008; Sato et al. 2011). For new assemblies, previously sequenced paired-end Illumina whole genome sequence reads were downloaded from NCBI’s Sequence Read Archive (Kjærbølling et al. 2020; Hatmaker et al. 2022)

Annotated protein-coding nucleotide sequences (CDS) for each nuclear genome were downloaded from JGI MycoCosm (Grigoriev et al. 2014; Kjærbølling et al. 2020) except for *A. sojae, A. flavus, and A. nomiae.* For *A. sojae,* strain-matched nuclear annotations were not available and thus this species was not included in phylogenetic inferences or any analyses based on nuclear genomic data. For *A. flavus* and *A. nomiae*, we used annotations from recently assembled genomes (Hatmaker et al., 2022), extracting the CDS regions. Strains and data sources are summarized in Table 1.

**Table 1:** Summary of sources of sequencing data. Reference mitochondrial genomes were used for *Aspergillus sojae*, *A. oryzae*, and *A. niger*. Raw paired-end whole genome sequencing reads were used for the remaining species.

| Species | SRA Accession/<br>NCBI Reference Sequence<br>GenBank | Source |
| --- | --- | --- |
| *Aspergillus flavus* | SRR18725159 | Hatmaker et al. 2022 |
| *Aspergillus transmontanensis* | SRR8398939 | Kjærbølling et al. 2020 |
| *Aspergillus arachidicola* | SRR8398876 | Kjærbølling et al. 2020 |
| *Aspergillus nomiae* | SRR19369914 | Hatmaker et al. 2022 |
| *Aspergillus parasiticus* | SRR8840397 | Kjærbølling et al. 2020 |
| *Aspergillus sergii* | SRR8840616 | Kjærbølling et al. 2020 |
| *Aspergillus sojae* | AP014506.1 | Sato et al. 2011 |
| *Aspergillus oryzae* | NC_008282.1 | Machida et al. 2005 |
| *Aspergillus minisclerotigenes* | SRR8398929 | Kjærbølling et al. 2020 |
| *Aspergillus caelatus* | SRR8840396 | Kjærbølling et al. 2020 |
| *Aspergillus pseudocaelatus* | SRR8840541 | Kjærbølling et al. 2020 |
| *Aspergillus pseudotamarii* | SRR8840579 | Kjærbølling et al. 2020 |
| *Aspergillus tamarii* | SRR8840604 | Kjærbølling et al. 2020 |
| *Aspergillus pseudonomiae* | SRR8840540 | Kjærbølling et al. 2020 |
| *Aspergillus bertholletius* | SRR8398880 | Kjærbølling et al. 2020 |
| *Aspergillus alliaceus* | SRR8396970 | Kjærbølling et al. 2020 |
| *Aspergillus coremiiformis* | SRR8398877 | Kjærbølling et al. 2020 |
| *Aspergillus leporis* | SRR8398928 | Kjærbølling et al. 2020 |
| *Aspergillus avenaceus* | SRR8839916 | Kjærbølling et al. 2020 |
| *Aspergillus novoparasiticus* | SRR8398934 | Kjærbølling et al. 2020 |
| *Aspergillus niger* | NC_007445.1 | Juhasz et al. 2008 |

### Mitochondrial Genome Assembly

Data from the whole genome sequence read files were extracted into usable format (FASTQ files) using SRA Toolkit v2.9.6-1 (Leinonen et al. 2011). Mitochondrial genomes were assembled from the raw reads of each species using the organelle genome assembler GetOrganelle v1.7.4.1 (Jin et al. 2020). Following the method of Hugaboom et al. (2021), we used the internal GetOrganelle fungal database (-F fungus_mt) and default parameter values for number of threads, extension, and k-mers to assemble the mitogenomes (Hugaboom et al. 2021). The complete mitochondrial genome for *Aspergillus fumigatus* SGAir0713 (GenBank accession: CM16889.1) was used as a reference for the seed database (parameter -s) for mitogenome assembly. Contigs generated for each *Aspergillus* species were circularized such that there was no overlap in the beginning and end of the mitochondrial genome sequence.

### Read Mapping

Read mapping to correct errors was carried out using Bowtie2 v2.3.4.1 (Langmead and Salzberg 2012) and SAMtools v1.6 (Li et al. 2009). Bowtie2 aligned the raw paired-end reads from each *Aspergillus* species against the corresponding circularized mitochondrial genome. Variants were identified using SAMtools. Read mapping was also visualized and variants identified using the Integrative Genomics Viewer (IGV) v2.9.4 (Robinson et al. 2017).

### Mitogenome Annotation

The rapid organellar genome annotation software GeSeq v2.03 (Tillich et al. 2017) was used to annotate the circularized mitochondrial genomes. In addition to the newly assembled mitogenomes, mitochondrial reference genomes for *A. oryzae* and *A. sojae* were also annotated using GeSeq. Gene names were adjusted, and translations were checked in accordance with the reference mitochondrial genomes of *A. flavus* TCM2014 (NC_026920.1), *A. oryzae* 3.042 (NC_018100.1), *A. parasiticus* (NC_041445.1), and *A. fumigatus* A1163 (NC_017016.1). Annotations were finalized following inspection of automated gene sequences using Geneious Prime v2021.1 (Kearse et al. 2012). OGDraw v1.1.1 (Greiner et al. 2019) was used for genome visualization.

### Multiple Sequence Alignment

Using MAFFT v7 (Katoh et al. 2019), single-gene multiple sequence alignment (MSA) files based on DNA nucleotide sequences were created for each of the 14 core mitogenes: cytochrome oxidase subunits 1, 2, and 3, NADH dehydrogenase subunits 1, 2, 3, 4, 4L, 5, and 6, ATP synthase subunits 6, 8, and 9, and cytochrome b. Gene nucleotide sequences corresponding to translated amino acid sequences for each gene were extracted from Geneious Prime v2021.1 (Kearse et al. 2012) sequence view and reverse complemented as necessary. The 14 individual MSA files were concatenated using SequenceMatrix v1.9 (Vaidya et al. 2011).

### Phylogenetic inference

To infer the evolutionary relationships within section *Flavi,* maximum likelihood phylogenies were constructed from both mitochondrial and nuclear data. The mitochondrial phylogeny was constructed from the MSA of 14 core concatenated mitogene nucleotide sequences files using RAxML v8.2.11 (Stamatakis 2014). The MSA was trimmed with ClipKIT v1.3.0 (Steenwyk et al. 2020) to retain parsimony-informative sites prior to construction of the phylogeny. *A. niger* (NC_007445.1) was used as the outgroup. We used 1,000 bootstrap replicates to evaluate robustness of inference. For the nuclear phylogeny, orthologous proteins in all species were identified using OrthoFinder v2.5.4 (Emms and Kelly 2019). MSAs for each of 2,422 orthologs were concatenated using the script catfasta2phyml.pl (https://github.com/nylander/catfasta2phyml). The maximum likelihood nuclear phylogeny was constructed with 1,000 replicates for bootstrapping using RAxML v8.2.11 (Stamatakis 2014) from the aligned orthologs shared among all the *Aspergillus* species in the study (including *A. niger*) except *A. sojae*, which did not have available sequencing data for nuclear genome assembly and annotation. The resulting consensus trees for both the mitochondrial and nuclear phylogenies were visualized using Geneious Prime v2020.1.2 (Kearse et al. 2012).

### Cluster Analysis

To compare patterns of synonymous codon usage bias between mitochondrial and nuclear genomes, hierarchical clustering of genome-level relative synonymous codon usage (RSCU) values was calculated and visualized using RStudio v. 2021.09.1. RSCU is a commonly used metric for codon usage bias that reflects the observed frequency of a particular codon divided by its expected frequency if all synonymous codons were used equally (Paul M Sharp and Li 1986). Genome-level RSCU values as well as RSCU values for each mitochondrial and nuclear gene were computed using DAMBE v7.3.5 (Xia 2017).

### Correspondence Analysis

To determine which codons drive differences in signatures of codon usage between nuclear and mitochondrial genes and between the mitogenomes of the 20 *Flavi* species, correspondence analyses were performed using gene-level RSCU values. Correspondence analysis was used for multivariate analysis because the RSCU values are interdependent—the RSCU values for one codon are inherently linked to the RSCU values of other synonymous codons—and thus not suited for principal component analysis. The correspondence analyses were carried out in RStudio v. 2021.09.1. using the packages ade4 v.1.7-19 (https://CRAN.R-project.org/package=ade4) and factoextra v.1.0.7 (https://CRAN.R-project.org/package=factoextra)

### Evaluation of mutational bias and codon usage

To evaluate the role of mutational bias in determining the observed patterns of codon usage bias in section *Flavi,* we plotted the effective number of codons (ENc) for each gene against their respective GC3 values, where GC3 is the GC content of the third codon position. ENc is often used to assess the non-uniformity of synonymous codon usage within individual genes (Wright 1990). Values range from 20 (extreme bias where only one codon is used per amino acid) to 61 (no bias). The ENc values for each gene were computed in DAMBE v.7.3.5 (Xia 2017). The resulting distribution was compared to the predicted neutral distribution proposed by dos Reis et al. (dos Reis et al. 2004) using the suggested parameters by computing the R^2^ values between the observed and expected ENc values.

### Evaluation of selection on codon usage

To compare the influence of translational selection on the codon usage bias of mitogenomes as compared to nuclear genomes, we calculated the S-value proposed by dos Reis et al. (2004) for each species. The S-value is the correlation between the tRNA adaptation index (stAI) and the confounded effects of selection on the codon usage of a gene as well as of other factors (e.g., mutation bias, genetic drift). Therefore, the S-value measures the proportion of the variance in codon bias that cannot be accounted for without invoking translational selection. Thus, the higher the S-value, the stronger the action of translational selection on the given set of genes.

To calculate the S-value, we first measured tRNA counts for each nuclear and mitochondrial genome using tRNAscan-SE 2.0 (Chan et al. 2021). These counts were used to calculate the species-specific value for each codon’s relative adaptiveness (wi) in stAIcalc, version 1.0 (Sabi et al. 2017). Exclusively mitochondrial tRNA counts were used to obtain wi values for mitogenomes, whereas exclusively nuclear genome tRNA counts were used for nuclear genomes. Taking the geometric mean of all wi values for the codons yielded the stAI of each gene. These stAI values were then used to calculate S-values for each mitochondrial and nuclear genome with the R package tAI.R, version 0.2 (https://github.com/mariodosreis/tai).

The statistical significance of each S-value was tested via a permutation test. 100 permutations were run such that each genome’s wi values were randomly assigned to codons, the tAI values recalculated for each gene, and the S-test run on that permutation. A genome’s observed S-value was considered statistically significant if it fell in the top 5% of the distribution formed by the 100 values obtained by the permutation analysis.

## Data Availability Statement

The newly assembled *Aspergillus* section *Flavi* mitogenomes from this study are available in GenBank under accession numbers ON833077, ON833078, ON833079, ON833081, ON833082, ON833083, ON833084, ON833085, ON833086, ON833087, ON833088, ON833089, ON833090, ON833091, ON833092, ON833093, and ON833094. Reannotations for previously assembled mitogenomes are available through figshare (https://figshare.com/s/ad503bfd4fc1b1914d10)

The SRA accession numbers for whole genome sequencing data used for mitogenome assembly are provided in Table 1. For previously assembled mitogenomes, the NCBI reference sequence

GenBank accession numbers are provided in place of SRA accession numbers.

Additional data are available through figshare (https://figshare.com/s/ad503bfd4fc1b1914d10).

## Results

### Genomic content varies greatly between nuclear and mitochondrial genomes

All mitogenomes were found to be small, circular DNA molecules with low GC content of 24.9-26.9% (Table 2). Fourteen core genes (cytochrome oxidase subunits 1, 2, and 3, NADH dehydrogenase subunits 1, 2, 3, 4, 4L, 5, and 6, ATP synthase subunits 6, 8, and 9, and cytochrome b of conserved order were contained in each annotated mitogenome (Figure 1). Additionally, a ribosomal protein S5 was found in all newly annotated *Flavi* genomes, and an intron encoded LAGLIDADG endonuclease was found in all mitogenomes except for *A. avenaceus* and *A. leporis.* Variations in mitogenome length are due to variations in intron number and length.

**Figure 1:**
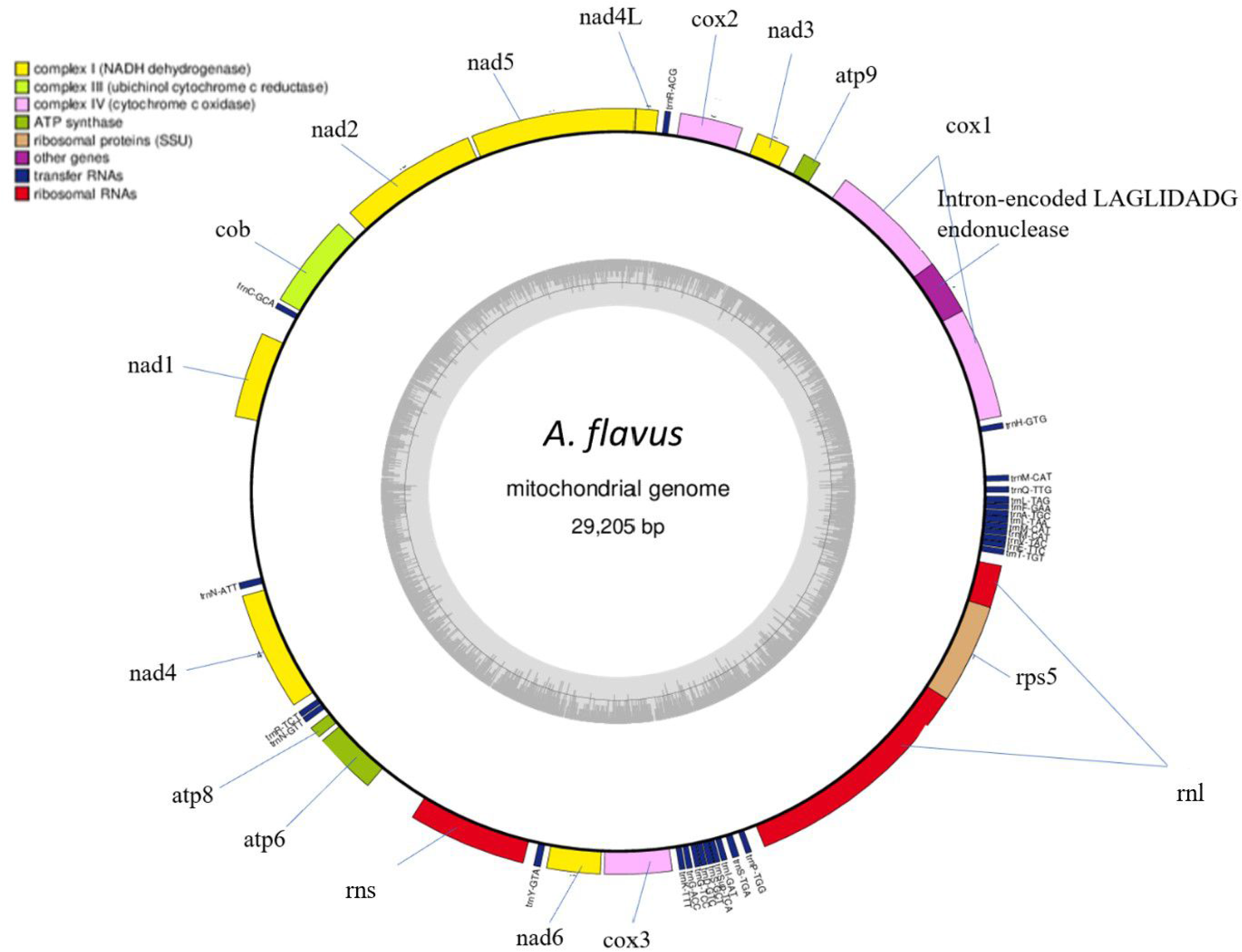
The typical *Aspergillus* section *Flavi* mitogenome is a circular DNA molecule of conserved gene content and order. Here, the circularized mitogenome of *Aspergillus flavus* 6333-EH-1is visualized. Each assembled section *Flavi* mitogenome shared a conserved set of 14 core mitochondrial genes, 2 rRNA genes, and 25-27 tRNA genes in the order pictured above. GC content (26.2% overall) is illustrated as the interior gray region.

**Table 2:** Mitochondrial and nuclear *Aspergillus* genomes differ greatly in size, genomic content, and GC bias. The summary above includes ranges of values from 20 *Aspergillus* section *Flavi* species’ mitochondrial and corresponding nuclear genomes.

|  | Genome length | rRNAs | tRNAs | CDS | GC% |
| --- | --- | --- | --- | --- | --- |
| Mitogenomes | 29.100-39.269 Kbp | 2 | 26 | 15-17 | 24.9-26.9% |
| Nuclear Genomes | 30,1001-40,900 Kbp | Undetermined | 228-272 | 9,078-14,216 | 43-48.8% |

Conversely, corresponding nuclear genomes are linear and have less extreme GC content biases ranging from 43.0-48.8% (Table 2). Nuclear genomes are roughly 1,000 times larger than their mitochondrial counterparts; while mitogenomes ranged from 29.100-39.269 Kbp, nuclear genomes ranged from 30,1001-40,900 Kbp. Of note, both nuclear and mitochondrial genomes house their own set of tRNAs (i.e., have their own tRNAome), although the tRNAome of nuclear genomes is roughly ten times larger than that of mitochondrial genomes. While nuclear genomes house 228-272 tRNAs, the mitogenomes encode a conserved set of 26 tRNAs. Importantly, each amino acid is represented by at least one tRNA in the conserved mitochondrial tRNAome. However, the codons GCC, GCU, CGG, CUC, CUU, CCC, CCU, UCC, UCG, UCU, ACC, ACU, GUC, GUU, UGG, UAG, and UAA could not be decoded by the set of unmodified mitochondrial tRNAs even when accounting for wobble base pairing.

### Mitochondrial and nuclear phylogenies are very similar

To understand how the evolutionary history of section *Flavi* informed by mitogenomes compares to that of nuclear genomes, a mitochondrial phylogeny was constructed using a concatenation of 14 core mitogene nucleotide sequences (Figure 2B). The resulting phylogeny displayed high bootstrap support. Despite minor topological differences from a well-supported nuclear phylogeny (Figure 2A) amongst more closely related species, the evolutionary relationships predicted by the mitochondrial phylogeny largely align with those predicted by the nuclear phylogeny. For instance, although *A. minisclerotigenes, A. sergii, A. flavus, A. arachidicola, A. parasiticus, and A. novoparasiticus* fall within the same clade in both nuclear and mitochondrial phylogenies, the predicted evolutionary relationships within this clade vary slightly. Evolutionary rate was found to be more rapid in mitochondrial genomes relative to nuclear genomes. For example, the evolutionary distance between *A. flavus* and *A. nomiae* was 0.086 substitutions per site in the nuclear phylogeny, but 0.244 substitutions per site in the mitochondrial phylogeny. Single-gene mitochondrial phylogenies differed in their topologies but exhibited low bootstrap support values, particularly for relationships among closely related species (Supplementary File S1).

**Figure 2:**
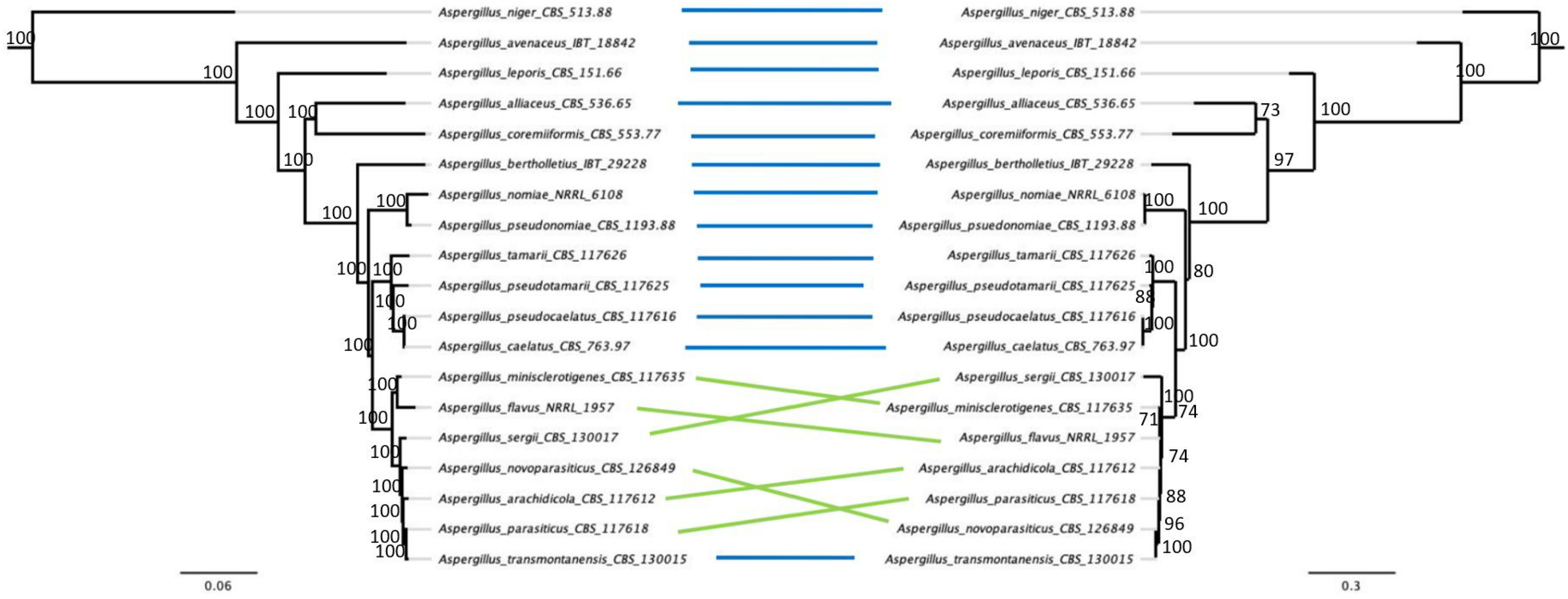
Phylogenies constructed from nuclear and mitochondrial data predict similar evolutionary relationships, with minor differences in inferred topology arising amongst more closely related species. A) Maximum likelihood phylogeny based on concatenation of 2,422 nuclear orthologs with bootstrap values from 1000 replicates. B) Maximum likelihood phylogeny based on concatenation of 14 core mitogene sequences with bootstrap values from 1000 replicates.

### Species groupings based on patterns of codon usage bias differ between mitochondrial and nuclear genomes

To compare similarities in interspecies patterns of CUB between mitochondrial and nuclear genomes, hierarchical clustering was performed using the net RSCU values of protein-coding regions of both nuclear (Figure 3A) and mitochondrial (Figure 3B) genomes. The cluster analyses predict different interspecies relationships depending on organelle of genomic origin. For example, the cluster dendrograms show that *A. oryzae* and *A. flavus* cluster together based on patterns of mitochondrial CUB, but group in completely different clusters based on nuclear CUB. This suggests that different pressures may govern CUB in mitochondrial genomes than in their nuclear counterparts.

**Figure 3:**
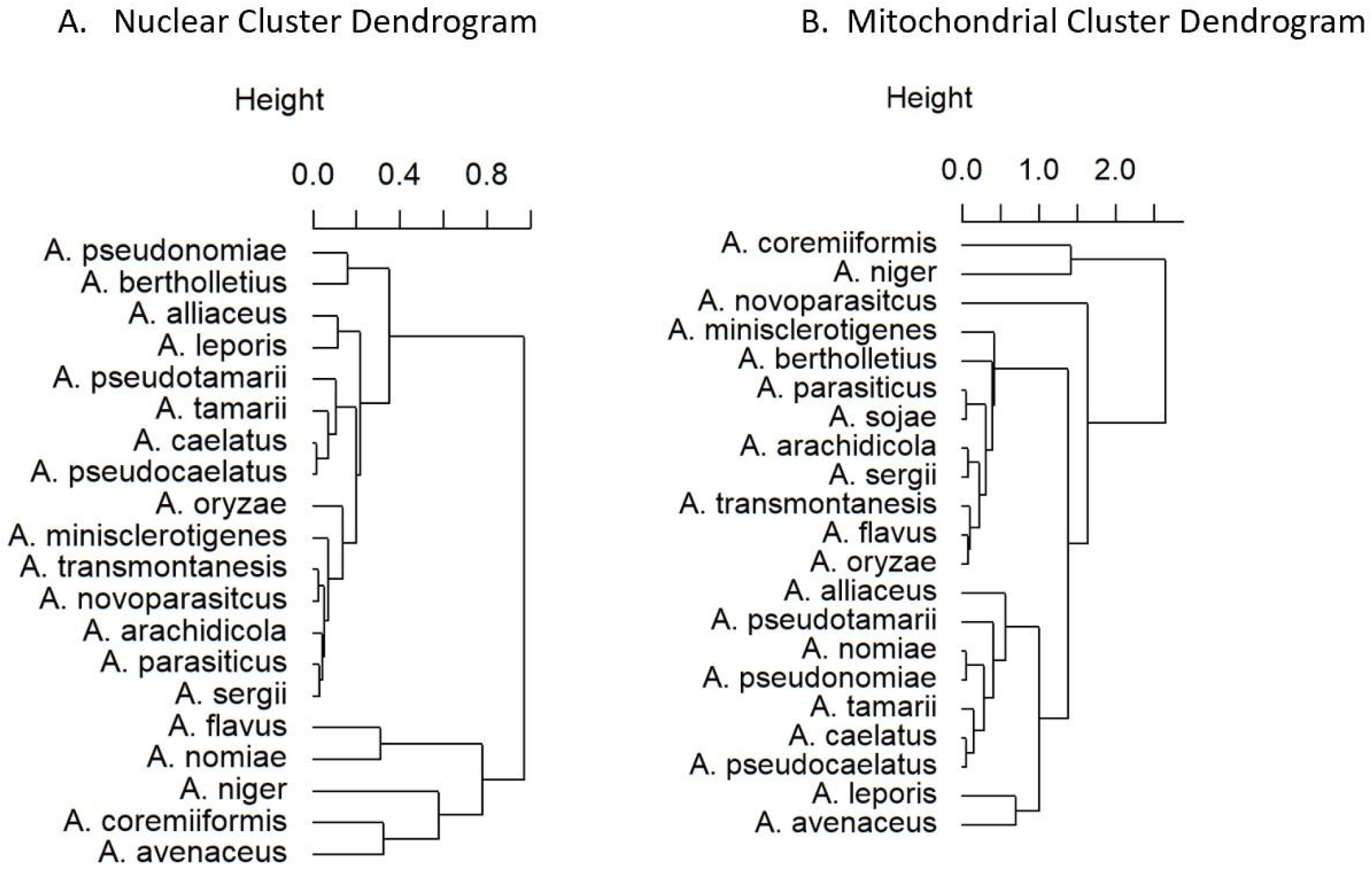
Hierarchical clustering analyses of relative synonymous codon usage (RSCU) values of mitochondrial and nuclear protein-coding regions demonstrate different species groupings based on mitochondrial and nuclear data. A) Cluster analysis based on net RSCU values of nuclear protein-coding genes B) Cluster analysis based on net RSCU values of mitochondrial protein-coding genes.

### Patterns of codon usage bias reflect whether genes are mitochondrial or nuclear in origin

To examine signatures of codon usage between nuclear and mitochondrial genes, RSCU values for each gene in each available genome were calculated. A correspondence analysis (CA) was then performed to determine which codons drive observed differences in codon usage patterns (Figure 4). The CA plot (Figure 4A) shows a distinct clustering of the majority of the mitogenes away from nuclear genes. This demonstrates that codon usage signatures depend more on whether genes are mitochondrial versus nuclear as opposed to whether genes belong to the same species. The factor map of codon contributions (Figure 4B) revealed that the first dimension explains 15.6% of observed variation between genes in the final plot. The second dimension explains 7% of observed variance. Examining dimensional contributions by codon reveals that the GC content of the third position drives separation along dimensions. Position along the first dimension (X-axis) is driven primarily by the usage of NNA versus NNC codons. The largest contributions along the X-axis come from the usage of AUA (isoleucine) and CCC (proline). RSCU values of greater than 1 indicate that a codon is overrepresented within a given synonymous codon group whereas RSCU values less than 1 indicate underrepresentation. The average RSCU of AUA and CCC are 1.6304 and 0.0449 in the mitochondria and 0.4525 and 1.0457 in the nucleus, respectively. Position along the second dimension (Y-axis) is driven primarily by differences in the usage of NNU versus NNG codons. The largest contributions along the Y-axis are from CCU (proline) versus GGG (glycine), ACG (threonine), and CCG (proline) combined. The average RSCU of CCU, GGG, ACG, and CCG are 2.7140, 0.0558, 0.0204, and 0.0484 in the mitochondria and 1.072, 0.7240, 0.8274, and 0.8920 in the nucleus, respectively.

**Figure 4:**
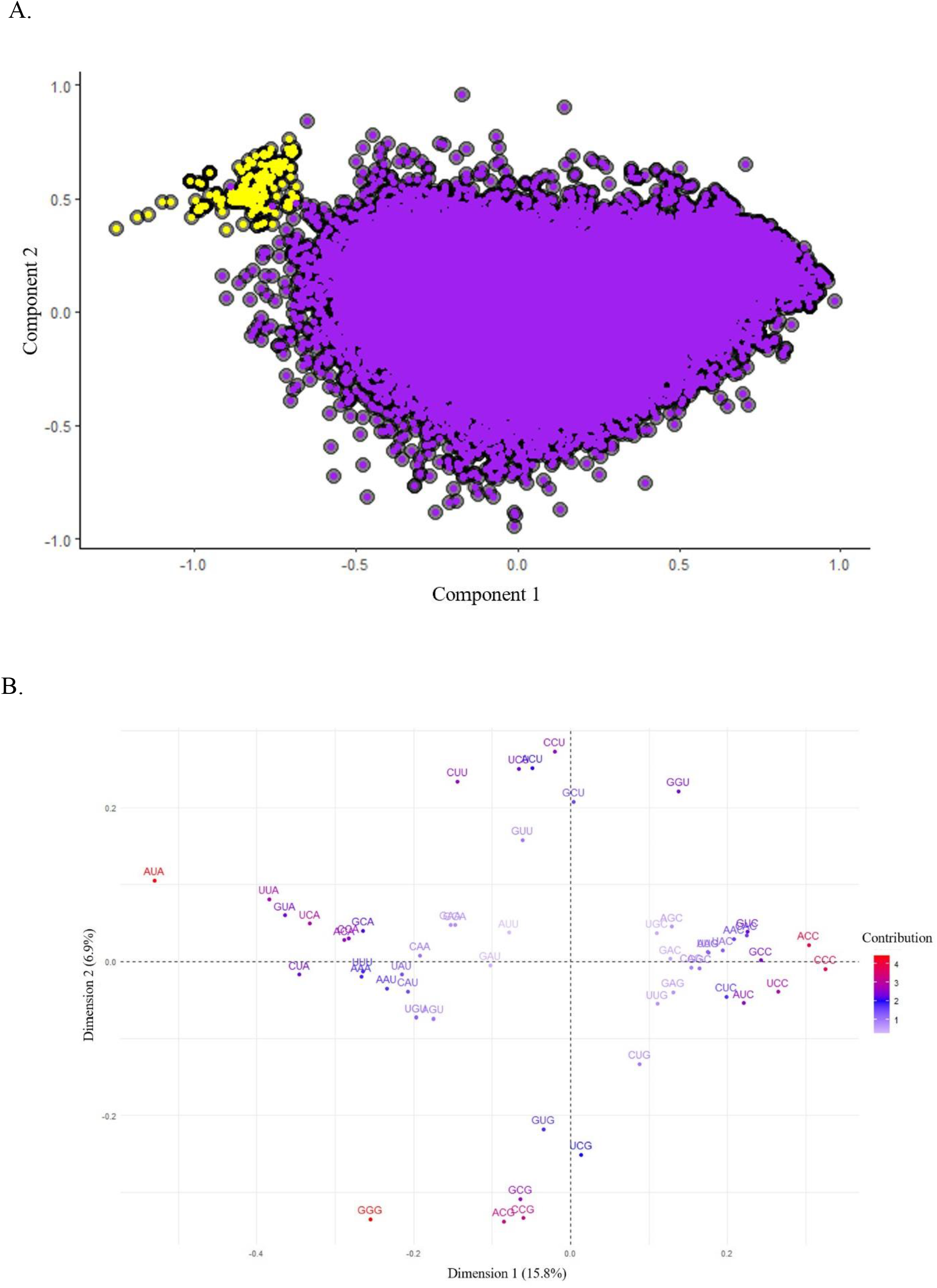
Correspondence Analysis based on relative synonymous codon usage values reveals that signatures of codon usage bias are more similar based on organelle of origin as opposed to species of origin. A) Correspondence analysis plot of all mitochondrial (yellow) and nuclear (purple) protein-coding genes for 20 *Aspergillus* section *Flavi* species. B) Factor map of codon contributions. Location of genes in correspondence analysis plot is driven largely by the GC content in the third position of synonymous codons used in the gene of interest.

A second CA was run using the RSCU values for each mitogene to determine which codons drives observed interspecies differences in codon usage patterns in mitogenomes (Figure 5). The CA plot shows distinct grouping based on gene identity as opposed to species of origin (Figure 5A). The factor map of codon contributions revealed that the first dimension explains 18.7% of observed variation in the final CA plot, while the second dimension accounts for 16.1% (Figure 5B). The *A. avenaceus atp8* gene is a clear outlier along both axes. The codons that contribute the most to this are ACC, UCC and CCG which are used at a frequency of 4, 4, 1.33 respectively in this gene. RSCU values of 4 indicate that only ACC (threonine) and UCC (serine)—none of the other synonymous codons within their respective families—are used in this gene. This degree of bias is expected given that threonine and serine occur only once and twice, respectively, in *A. avenaceus atp8*.

**Figure 5:**
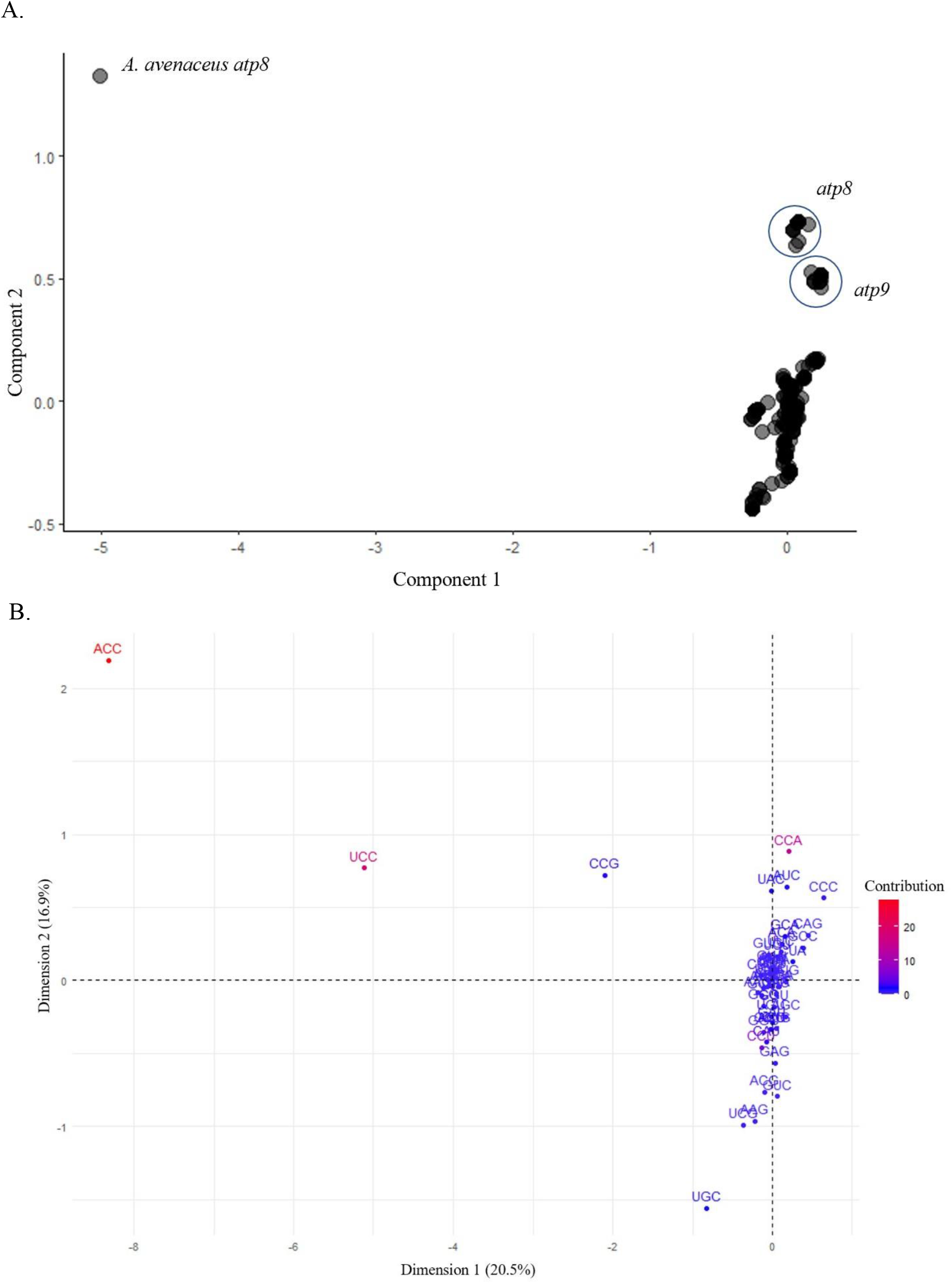
Correspondence Analysis based on relative synonymous codon usage values in mitogenes reveals that signatures of codon usage bias are more similar based on gene identity as opposed to species of origin. A) Correspondence analysis plot of all mitochondrial protein-coding genes for 20 *Aspergillus* section *Flavi* species. Labels correspond to gene identity B) Factor map of codon contributions

### Deviation of gene-level codon usage from neutral expectation varies based on whether genes are of nuclear or mitochondrial origin

To assess the role of mutational bias across all mitochondrial and nuclear genes, we examined the relationship between the ENc of each gene and its GC3 content by comparing observed ENc values to the expected relationship between ENC and GC3 content if codon usage was influenced by neutral mutational bias alone. We tested the fit to the neutral expectation of the complete dataset of all species’ combined nuclear and mitochondrial gene datasets as well as all nuclear genes and all mitogenes separately by calculating the R^2^ value. For all 20 species, combined nuclear and mitochondrial datasets yielded R^2^ values greater than 0.5, suggesting that codon usage in these species can be partially explained by neutral mutational bias (Supplemental File S2). Furthermore, patterns of deviation from the neutral expectation were highly similar between species (Supplemental File S2). However, when nuclear and mitochondrial genes were analyzed separately, nuclear genes had an R^2^ value of 0.598, whereas mitochondrial genes had an R^2^ value of 0.211 (Figure 6). This suggests that, although codon usage in nuclear genomes can be partially explained by neutral mutational bias, mutational bias does not fully account for the codon bias in mitochondrial genomes.

**Figure 6:**
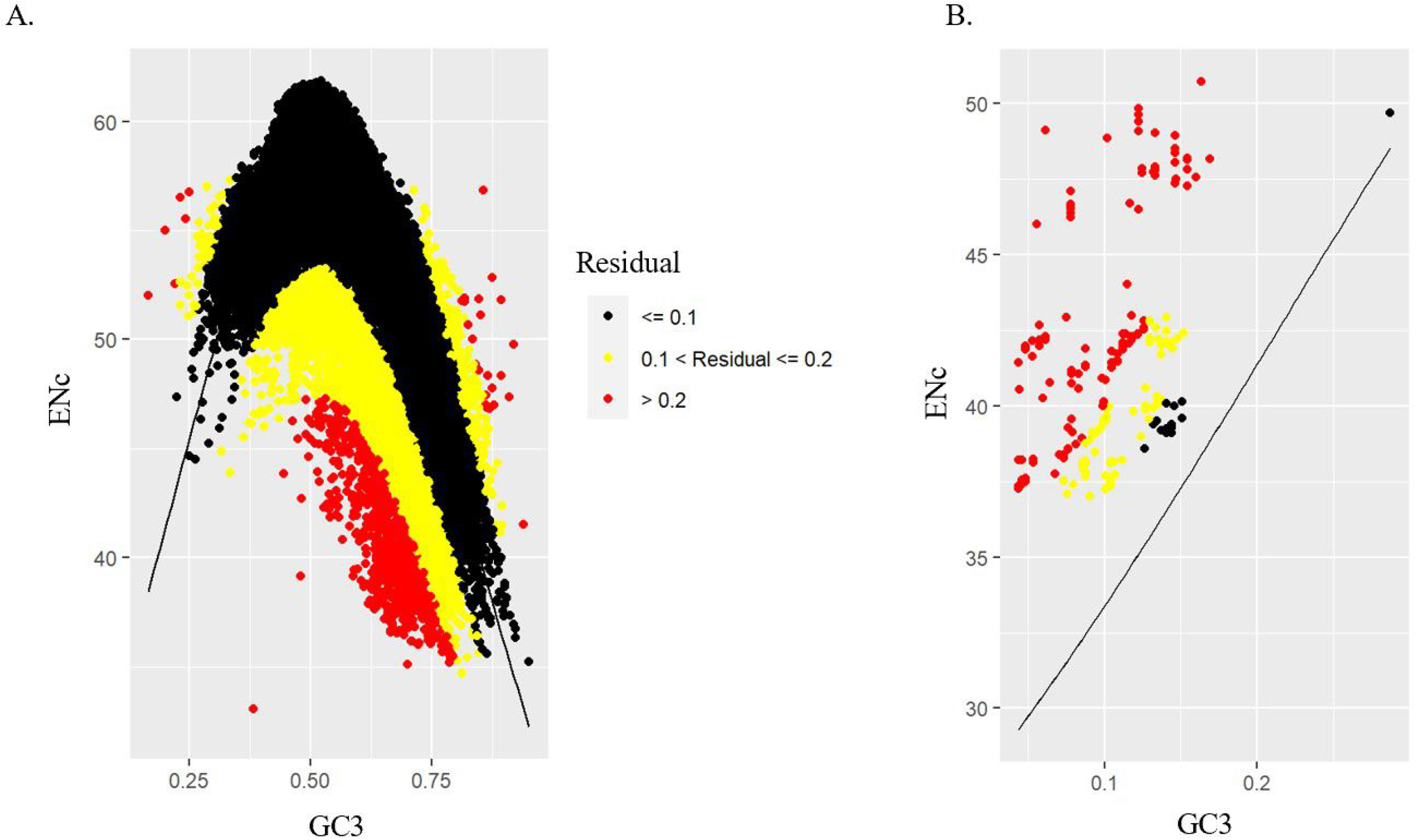
Most signatures of codon usage bias in mitochondrial and nuclear genes in *Aspergillus* section *Flavi* deviate from the expected codon usage bias under mutation pressure alone. A) ENc-GC3 plot for all nuclear protein-coding genes of 20 *Aspergillus* section Flavi genes plotted against the predicted neutral distribution. R^2^ value of 0.598 indicates moderate fit to neutral expectation. B) ENc-GC3 plot for all protein-coding mitogenes. R^2^ value of 0.211 indicates poor fit to neutral expectation.

### Codon usage in nuclear genomes, but not mitogenomes, is under translational selection

To test if translational selection could account for the observed deviations of CUB from the neutral expectation, we calculated the S-values for each mitochondrial and nuclear genome. Of the 20 *Flavi* species tested, mitogenome S-values ranged from -0.103 to 0.392 with a median value of 0.162 and mean value of 0.137 (Figure 7A). However, no species’ mitogenomes had S-values that were found to be significant in the permutation test. In contrast, nuclear genome S-values ranged from 0.269 to 0.502, with a median value of 0.432 and a mean value of 0.427 (Figure 7B). The S-value of *A. novoparasiticus* (S = 0.269) was calculated using the package tAI.R (https://github.com/mariodosreis/tai/blob/master/R/tAI.R). This was done as the original calculation with stAI calc created issues with file merging. All nuclear S-values were found to be significant in the permutation test, suggesting that *Flavi* nuclear genomes are under moderate levels of translational selection.

**Figure 7:**
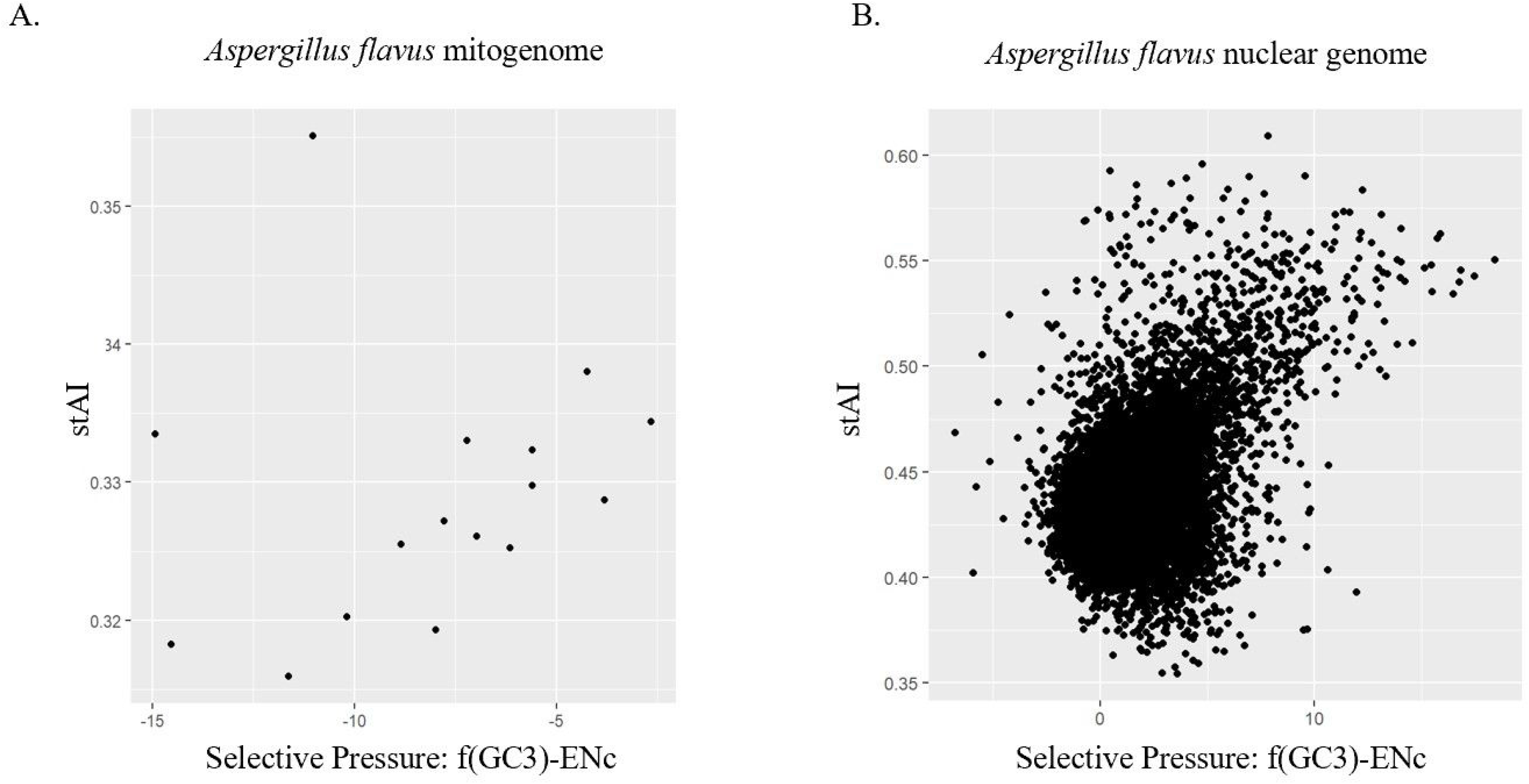
Section *Flavi* mitogenomes are not under significant translational selection on codon usage bias, but nuclear genomes display moderate translational selection. Plots of stAI against selective pressure for all protein-coding genes of *Aspergillus flavus* A) Mitogenes only. Example of insignificant translational selection on S-test (S=0.191). B) Nuclear genes only. Example of moderate translational selection on S-test (S=0.454).

## Discussion

In this study, we compared the evolution of mitochondrial and nuclear genomes within *Aspergillus* section *Flavi*. We assembled and annotated the mitogenomes of 18 *Flavi* species and reannotated two previously assembled reference mitogenomes. We then used phylogenetic analyses to compare phylogenies derived from nuclear versus mitochondrial data. Finally, we examined the patterns of and forces underlying CUB in nuclear and mitochondrial genomes.

The newly assembled mitogenomes are comparable in gene content and size to previously published *Aspergillus* mitogenomes. At 29.10 kb to 39.27 kb, the range of *Aspergillus* section *Flavi* mitogenome length falls within the lower range of published fungal mitogenomes, which vary in size from 12.06 kb to 235.85 kb (Joardar et al. 2012; Zhang et al. 2020). GC content was consistent with low percentages observed in other *Aspergillus* and related fungal species (Machida et al. 2005; Sato et al. 2011; Joardar et al. 2012; Zhao et al. 2012; Yan et al. 2016; Park et al. 2019a; Park et al. 2019b; Park et al. 2020; Hugaboom et al. 2021). The mitogenomic content and gene order were highly conserved in the 20 *Flavi* species analyzed, and all mitogenomes examined contained 14 core mitochondrial genes. As in previous studies, these core genes showed high levels of sequence similarity and conservation of gene order (Joardar et al. 2012; Hugaboom et al. 2021). Fungal mitogenomes are also known to contain accessory genes in addition to the core set of 14. The presence of the two accessory genes—an intron encoded LAGLIDADG endonuclease and the ribosomal protein S5—in most of the species analyzed is also consistent with existing *Flavi* annotations (Joardar et al. 2012; Hugaboom et al. 2021). The order of these accessory genes was also highly conserved. Of note, the mitogenomes contained their own set of 26 tRNAs separate from the nuclear-encoded set of tRNAs. In analyses of CUB, the mitochondrial tRNAs were used to determine if mitochondrial CUB patterns had been optimized to the mitochondrial tRNA pool.

In comparing the topologies and evolutionary rates predicted by phylogenies derived from nuclear and mitochondrial data, we found that their inferred evolutionary histories were similar. The high degree of congruence in the two phylogenies suggests potential coevolution of mitochondrial and nuclear genes. The minor disagreements between the two phylogenies may be explained by phenomena that occur uniquely in the mitochondria. For example, fungal mitochondria are uniparentally inherited (Santamaria et al. 2009). Additionally, fungal species can undergo interspecific hybridization (Giordano et al. 2018). In this process, the mitochondria of one species may be inherited by the other. More rarely, mitochondrial recombination events with repeated backcrossing can lead to introgression (Giordano et al. 2018). Interspecific introgression and recombination occur in fungal nuclei as well. Thus, the two phylogenies may differ due to introgression or recombination occurring in the mitochondria or nuclei of section *Flavi.* Alternatively, however, topological differences could arise due to sampling error, as mitochondrial genes contain few sites relative to nuclear genes, for example.

Cluster analyses based on net RSCU values demonstrated that interspecies similarities in patterns of CUB differ between nuclear and mitochondrial genomes. Despite some parallels in predicted grouping—for example, the grouping of *A, caelatus* and *A. pseudocaelatus* in both dendrograms —mitochondrial and nuclear cluster analyses displayed groupings largely inconsistent with each other. It is important to note that all the species included in this study have similar mean values of codon usage metrics (ENc, GC content, and GC3s) within the nuclear and mitochondrial genome. Thus, well-resolved interspecies relationships are unlikely to be had based on codon usage indices alone. Alternatively, the observed incongruence may reflect different pressures governing CUB in mitochondrial compared to nuclear genomes.

Examination of codon usage patterns using correspondence analyses showed that differential usage of certain codons drives observable differences in signatures of CUB between mitochondrial and nuclear genes and between gene type in mitogenes. Differential usage of specific codons between nuclear and mitochondrial genomes appears to rely heavily on the GC content of the third position of synonymous codons. This pattern aligns with overall GC content of the genomes. For example, the use of the codon AUA contributes to the placement of the mitogenomes in quadrant II of the final correspondence analysis plot, where mitogenes tend to cluster, while the use of AUC contributes to the placement of the nuclear genes in quadrants I and IV. The average RSCU values of AUA and AUC are 1.6304 and 0.1782 in the mitochondria and 0.4525 and 1.4760 in the nucleus, respectively. Both of these codons code for isoleucine, yet mitogenes are enriched for the AUA codon and nuclear genes for the AUC codon, as would be expected based on the differences in GC content between the two genomes. The separation of mitogenes is also dependent on the GC content of the third position. Figure 5 shows that, while the use of most codons is similar amongst all mitogenes, the occurrence of a rare G- or C-ending codon drives separation based on CUB patterns. This is especially clear in the case of the outlying *A. avenaceus atp8* gene in Figure 5A, which is driven by the higher use of codons ACC, UCC, and CCG. Despite a high degree of sequence conservation with the other 19 *atp8* nucleotide sequences (Supplementary File S3), the change in a small number of nucleotides at third codon positions results in a large visible separation in the correspondence analysis plot (Figure 5A). This effect is amplified due to the short, highly conserved nature of the *atp8* gene sequences. Overall, we found that gene-level RSCU values allow for observable differences in CUB pattern based on the organelle of genomic origin and mitogene identity.

We also sought to determine the relative importance of neutral processes and natural selection on shaping CUB in mitochondrial and nuclear genomes. Based on ENc-GC3 plots, most mitogenes fell at least 20% from the neutral expectation, while most nuclear genomes fell within 10% of the neutral expectation. These results reinforce previous findings that CUB varies at the gene-level within a species (Sharp et al. 1988; L et al. 2004; LaBella et al. 2019). Of note, studies have shown that greater divergence from the neutral expectation is moderately associated with increased expression (Tsankov et al. 2010). Future avenues may examine the association between the large residuals from the neutral expectation and expression levels of mitogenes.

The moderate to poor fit to the neutral expectation for nuclear and mitochondrial genes, respectively, suggests that mutational bias alone cannot account for the observed patterns in codon usage bias. By using the S-test to test for the influence of translational selection, we found that gene-level codon usage in mitochondrial genomes could not be significantly distinguished from neutral mutational bias in section *Flavi*, while translational selection acts moderately on codon usage bias in nuclear genomes. The lack of significant translational selection on mitogenomes is unsurprising, given their extreme GC bias and small size. This may be a manifestation of mtDNA evolving clonally with limited ability to recombine; thus, CUB is more likely to reflect mutation bias and drift rather than selection. Additionally, when genome size is small, it is hypothesized that low tRNA redundancy limits the ability of selection to act on CUB (dos Reis et al. 2004). Of note, S-value calculation for mitogenes was limited to a dataset of 16 genes. Visual inspection of the data used to determine the S-values suggests a general positive correlation between selective pressure and codon usage – which would suggest translational selection on codon usage – that is obscured by a couple outlier genes (Figure 7A). This observation in combination with the highly variable codon usage between mitochondrial genes suggests that the balance between selective and neutral forces on mitochondrial codon usage may vary greatly between mitogenes. Finally, the final S-value calculations for mitogenomes were based solely on the mitochondrial tRNA counts derived from genomic sequences and not experimental tRNA abundances. In fact, our analysis suggests that additional tRNA dynamics, such as modification or importation, may be at work in *Aspergillus* mitochondria.

Computational analysis of codon usage and tRNA composition in *Aspergillus* mitogenomes suggests that there is a significant gap in our knowledge of tRNA dynamics within these organelles. It is known that mitochondria can employ diverse strategies to obtain a complete and functional set of tRNAs. Some organisms such as the fungus *Saccharomyces cerevisiae* encode a complete set of tRNA genes within their mitochondria (Salinas-Giegé et al. 2015) while others require the importation of nuclear tRNAs into the mitochondria (Alfonzo and Söll 2009). Our analysis demonstrates that even when wobble base pairing is accounted for, 17 codons cannot be decoded by the mitochondrial tRNAome in the absence of tRNA modification (Supplementary File S4). This suggests that tRNA import or modification may be occurring in *Aspergillus* mitochondria. Additionally, mitochondrial codon usage is not consistently biased towards codons matching the mitochondrial tRNAome. For example, the codon GCA (alanine), which can be decoded by a mitochondrial tRNA, has an average RSCU value of 1.4222 in mitogenomes, whereas the codon GCU (also alanine) has an average RSCU value of 2.3423 even though the mitochondrial tRNAome is unable to decode this codon. The preference for GCU codons suggests the importation or modification of a tRNA capable of decoding this codon. Finally, the *Aspergillus* mitochondrial tRNAs fit the wobble versatility hypothesis for each codon family, with the exception of CGN (arginine), UGR (tryptophan), and AUR (methionine), a finding that is consistent with previous investigation of the wobble nucleotide position in fungal mitogenomes (Supplemental File S5) (Carullo and Xia 2008). That is, the anticodons of the mitochondrial tRNAome have nucleotides at the wobble site that maximize versatility in wobble base pairing as opposed to maximizing Watson-Crick base pairing with the most frequently used codon within each synonymous codon family. Improving our understanding of *Aspergillus* mitochondrial tRNA dynamics will not only allow us to better understand translational dynamics within the organelle but recent work has suggested that mitochondrial tRNAs may play a role in antifungal response (Colabardini et al. 2022).

Despite a limited understanding of tRNA dynamics within *Aspergillus* mitochondria our results are consistent with the limited role of translation selection in shaping general patterns of mitochondrial codon usage in other species including budding yeasts, plants, and animals (Kamatani and Yamamoto 2007; Jia and Higgs 2008; Zhou and Li 2009). As with previous work, we also noted a few specific codons (proline codons) and genes (*atp8*) with increased biases that may be related to factors such as wobble-decoding or tRNA abundance.

In summary, analysis of mitochondrial and nuclear genome data from *Aspergillus* section *Flavi* revealed that both genomes are largely phylogenetically congruent and that the pattern and evolutionary forces shaping CUB differ between the mitochondrial and nuclear genomes. These evolutionary analyses, coupled with the generation of mitogenome assemblies for 18 section *Flavi* species, contribute to our understanding of genome evolution in the genus *Aspergillus*.

## Acknowledgements

We thank members of the Rokas Laboratory for their support and feedback. This work was performed in part using resources contained within the Advanced Computing Center for research and Education at Vanderbilt University in Nashville, TN.

## Supplementary Materials

Available through figshare (https://figshare.com/s/ad503bfd4fc1b1914d10).

